# Building capacity among caregivers for managing children with Type 1 Diabetes Mellitus through a structured Interprofessional training program

**DOI:** 10.1101/2024.10.31.621438

**Authors:** Ayesha Juhi, Saleena Ummer Valladath, Himel Mondal, Rajan Kumar, Sangita Bhattacharya, Santanu Nath, Anupkumar Dhanvijay, Mukhayprana Prabhu

## Abstract

**Background:** Effective management of type 1 diabetes mellitus (T1DM) in children necessitates not only medical care but also substantial involvement from caregivers, who often encounter psychological, financial, and emotional challenges. This study aimed to develop and evaluate a structured educational program for caregivers of children with T1DM, facilitated by an interprofessional team.

**Methods:** This mixed-method study, conducted from May 15 to August 15, 2023, comprised two parts. First, a structured educational module was developed based on a needs assessment from semi-structured interviews with 20 caregivers. The module was created in English, Hindi, and Khortha (the local language of Deoghar, Jharkhand, India) and validated by a panel of 10 experts (CVI = 0.83). It was delivered using audio-visual aids, live demonstrations, posters, and booklets. The program’s effectiveness was evaluated through a pre-post-test questionnaire.

**Results:** The interviews revealed themes such as misunderstandings of treatment plans, distress from frequent insulin administration, fear of glucose monitoring, financial burdens, and psychological stress. Data from 68 of the 75 caregivers who participated in the educational program were analyzed. The Wilcoxon signed-rank test indicated a significant improvement in caregiver knowledge and management practices (P < 0.001), with a large effect size (Cohen’s d = 3.2).

**Conclusion:** This study demonstrates that structured educational interventions can significantly enhance caregivers’ knowledge and practices in managing T1DM. Multidisciplinary education and support are crucial for addressing the emotional and financial burdens caregivers face, ultimately improving the well-being of both caregivers and children.

## INTRODUCTION

In 2021, approximately 1.5 million individuals under the age of 20 worldwide were affected by Type 1 diabetes mellitus(T1DM).^1^ It is an autoimmune disease in children, often presents suddenly and without much warning. By the time of diagnosis, one-third of affected children may develop life-threatening diabetic ketoacidosis.^2^ Ensuring high-quality care for T1DM, particularly during childhood and adolescence, is essential for achieving optimal metabolic control and long-term health outcomes. Parents/caregivers play a critical role in diabetes management for children with T1DM. Indeed, children, adolescents, and their families face significant psychological distress upon the diagnosis of T1DM and endure a challenging and lengthy adjustment process. Parental adaptation plays a crucial role, either negatively or positively impacting a child’s ability to cope with T1DM.^3-4^ Diabetes care should prioritize a model that focuses on placing child and their parents/caregivers at the centre of the care approach. Managing diabetes requires navigating unpredictable activity, routines, and dietary patterns as children develop across physiological, cognitive, behavioural, social, and emotional dimensions.^5^ Caregivers of children with T1D often face elevated levels of stress, depression, and anxiety.^6^

Given the complexity of T1DM management, there is a critical need for interprofessional collaboration to educate caregivers effectively in supporting children with T1DM. This collaboration is essential for creating awareness and managing the daily tasks such as assessing blood glucose levels, administering insulin, regulating food intake, managing physical activity, providing psychological support, and preventing complications associated with T1DM.

Providing caregivers with comprehensive education on disease management and empowers them to effectively manage their child’s T1DM, this holistic approach ensures the child’s physical health, emotional well-being, and overall quality of life are optimized, while reducing the stress and burden on caregivers.

Hence this mandates the need of the present study. This study was taken up with the objectives to develop a structured educational program on disease management for caregivers of children with T1DM involving an interprofessional team and evaluate its effectiveness through a pre-post-test analysis.

## MATERIALS AND METHODS

### Type and settings

This mixed-method study was conducted with caregivers of children with type 1 diabetes mellitus (T1DM) from May 15 to August 15, 2023. The study received approval from both the Institutional Research Committee and the Institutional Ethical Committee. Caregivers were recruited from the pediatric outpatient department of a tertiary care hospital and community health centers in nearby villages. The study consisted of two parts: the first involved a needs assessment through in-depth interviews and the development and validation of educational materials, while the second involved the implementation of the educational intervention and assessment of its effectiveness. A brief overview of the study process is illustrated in Figure 1.

**Figure 1:**
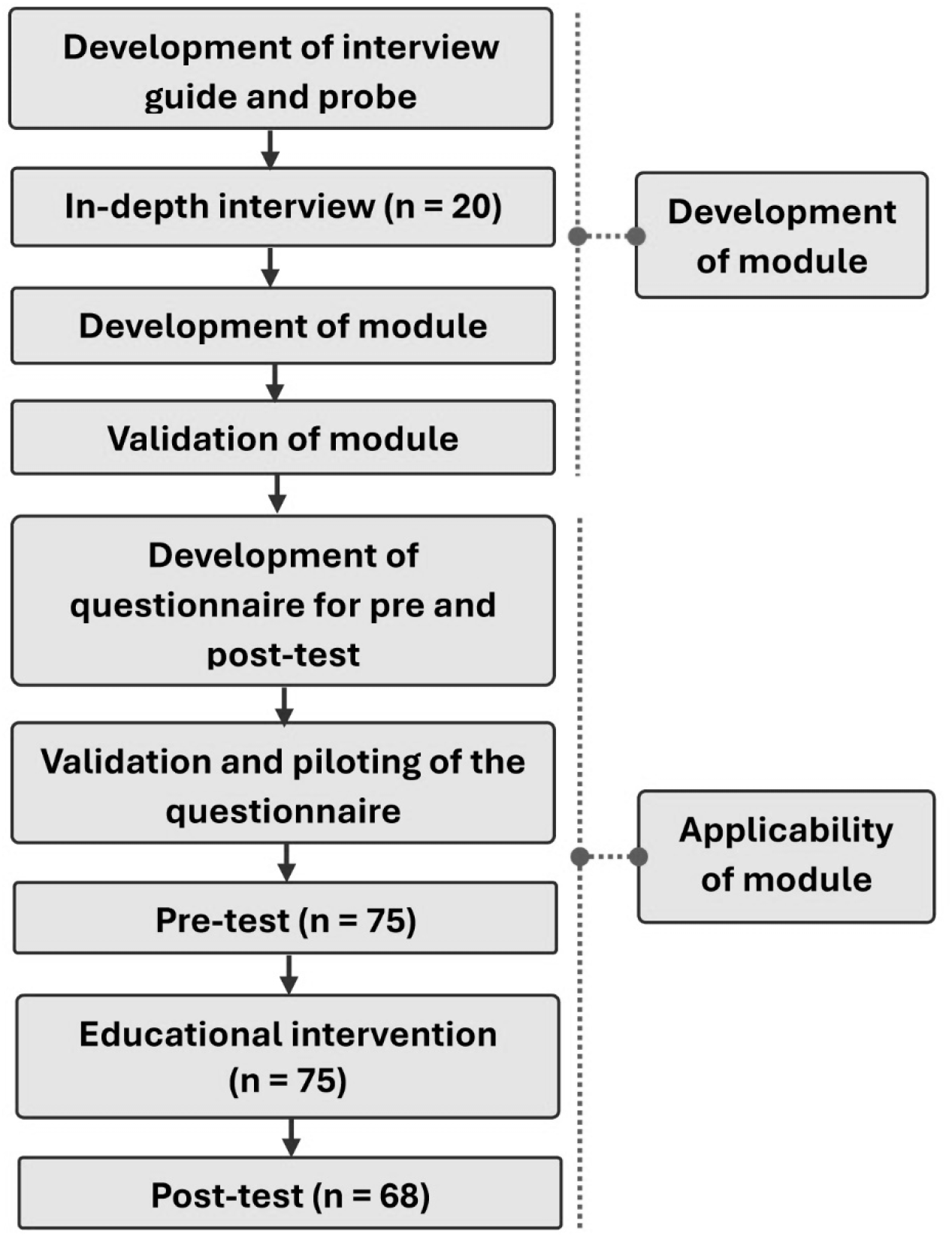
Overall study procedure.

### In-depth Interviews

A needs assessment was conducted using semi-structured interviews with 20 caregivers, each lasting 30-45 minutes. These interviews explored caregivers’ experiences, knowledge, awareness, and practices in managing T1DM, as well as the psychosocial and economic challenges they face. The interviews were conducted by an experienced interviewer using a pre-designed guide and probes, with prior experience in health-related studies.

### Generation of Themes

The interviews were recorded and manually transcribed by an independent individual proficient in English, Hindi, and the local language. The transcriptions were thematically analyzed by two authors (AJ and HM), who reached a consensus on the final themes. Identified themes included understanding T1DM, insulin management, dietary practices, physical activity, psychological support, and handling emergencies. A comprehensive educational module was developed to address these specific needs and challenges.

### Development of Educational Material

The educational materials were meticulously crafted based on the themes identified in the interviews. A multidisciplinary team of experts contributed to the content development, which included three paediatricians, two endocrinologists, two clinical dietitians, clinical biochemist, a psychiatrist, a family medicine specialist, a health promotion expert, and an exercise and yoga therapist. The materials were produced in English, Hindi, and Khortha to ensure cultural relevance and accessibility for local caregivers.

### Validation of the Module

The module underwent validation by a panel of three experts, including pediatricians and a psychiatrist, who not been involved in the material’s creation. The module achieved high face validity (>80%) and content validity (CVI = 0.825).

### Pre-test and Post-test Questionnaire Development

An 11-item questionnaire was developed by the interprofessional team to evaluate the educational module’s effectiveness. This questionnaire, designed to assess awareness and knowledge of disease management, was validated for content and pre-tested on 10 participants. The complete questionnaire is provided as Annexure 1.

### Intervention (Training of Caregivers)

Before the intervention, participants completed the 11-item questionnaire. An expert surveyor collected responses, after which a fixed team of four interprofessional healthcare members (one paediatrician, one nutritionist, one nurse, and one exercise therapist) conducted training sessions for groups of five caregivers, each lasting approximately two hours. Content was delivered through audio-visual aids and live demonstrations on insulin titration, diet, and physical exercise. Participants also received educational materials, including posters and booklets with pictorial content. Individual follow-up sessions were offered to those needing further assistance or clarification on disease management. Six weeks later, caregivers completed the same questionnaire for follow-up assessment.

### Added Intervention

The participants from the in-depth interviews (n = 20) underwent similar educational intervention sessions, although their data were not included in the analysis.

### Statistical Analysis

Data were expressed as numbers and percentages. Categorical data were compared using the Chi-square test. Since the pre- and post-test scores were not normally distributed, the Wilcoxon signed-rank test was employed for comparison. Statistical analysis was conducted using GraphPad Prism 9.5.0 (GraphPad Software, USA), with a significance level set at p < 0.05.

## RESULTS

The needs assessment was conducted with 20 caregivers (Table 1) who participated in in-depth interviews. Major themes and subthemes identified included misunderstandings of the treatment plan, difficulties in titrating the correct dosage of insulin, confusion about appropriate food choices, and uncertainty about when to seek emergency medical help. Caregivers expressed distress over administering frequent insulin shots, which caused unease for the child and led to fears of multiple pricks, missed doses, and avoidance of social gatherings. Another significant concern was the fear associated with glucose monitoring; many children cried at the sight of the lancet, and caregivers were distressed by their child’s painful, blackened fingertips.

**Table 1:** Characteristics of caregivers.

| **Table 1: Characteristics of caregivers** |  |  |  |
| --- | --- | --- | --- |
| **Attribute** | **Category** | **Interview group (n = 20)** | **Intervention group (n = 68)** |
| Age (years) | 18–24 | 2 | 6 |
|  | 25–34 | 3 | 19 |
|  | 35–44 | 9 | 28 |
|  | >44 | 6 | 15 |
| Sex | Males | 7 | 31 |
|  | Females | 13 | 37 |
| Relation | Mother | 9 | 22 |
|  | Father | 6 | 19 |
|  | Sibling | 2 | 8 |
|  | Grandparent | 0 | 9 |
|  | School attender | 0 | 2 |
|  | Uncle | 2 | 3 |
|  | Aunt | 1 | 5 |
| Education level | Education deprived | 5 | 12 |
|  | <10<sup>th</sup> standard | 2 | 22 |
|  | 10<sup>th</sup> - 12<sup>th</sup> standard | 4 | 19 |
|  | Graduate | 5 | 11 |
|  | Post graduate and above | 4 | 4 |
| Family income (INR/month) | <10000 | 5 | 12 |
|  | 10001- 20000 | 4 | 17 |
|  | 20001 – 30000 | 3 | 19 |
|  | 30001- 40000 | 5 | 10 |
|  | 40001- 50000 | 2 | 8 |
|  | >50000 | 1 | 2 |

Financial burdens emerged as a major issue, with caregivers struggling to meet their child’s special needs, afford monthly treatment costs, and pay bills post-hospitalization. There was a clear call for government assistance, such as discounts or free insulin. Psychological stress was prevalent among caregivers, as they witnessed their child’s inability to engage in physical activities with peers, faced social isolation, feared potential emergencies at school, and reconsidered having more children due to concerns about T1DM.

A total of 75 participants were recruited for the intervention. All completed the pre-test; however, 7 caregivers dropped out by the 6-week follow-up, resulting in data from 68 caregivers included in the analysis. Demographic details of the study participants are presented in Table 1. The Pre-test and Post-test data were summarized as mean±standard deviation (SD). The mean Pre-test score was 14.26±3.34, while the post-test score was 23.17±2.28. A statistically significant increase in post-test scores was observed (Wilcoxon signed-rank test, P < 0.001), as illustrated in Figure 2. The effect size was large (Cohen’s d = 3.2).

**Figure 2:**
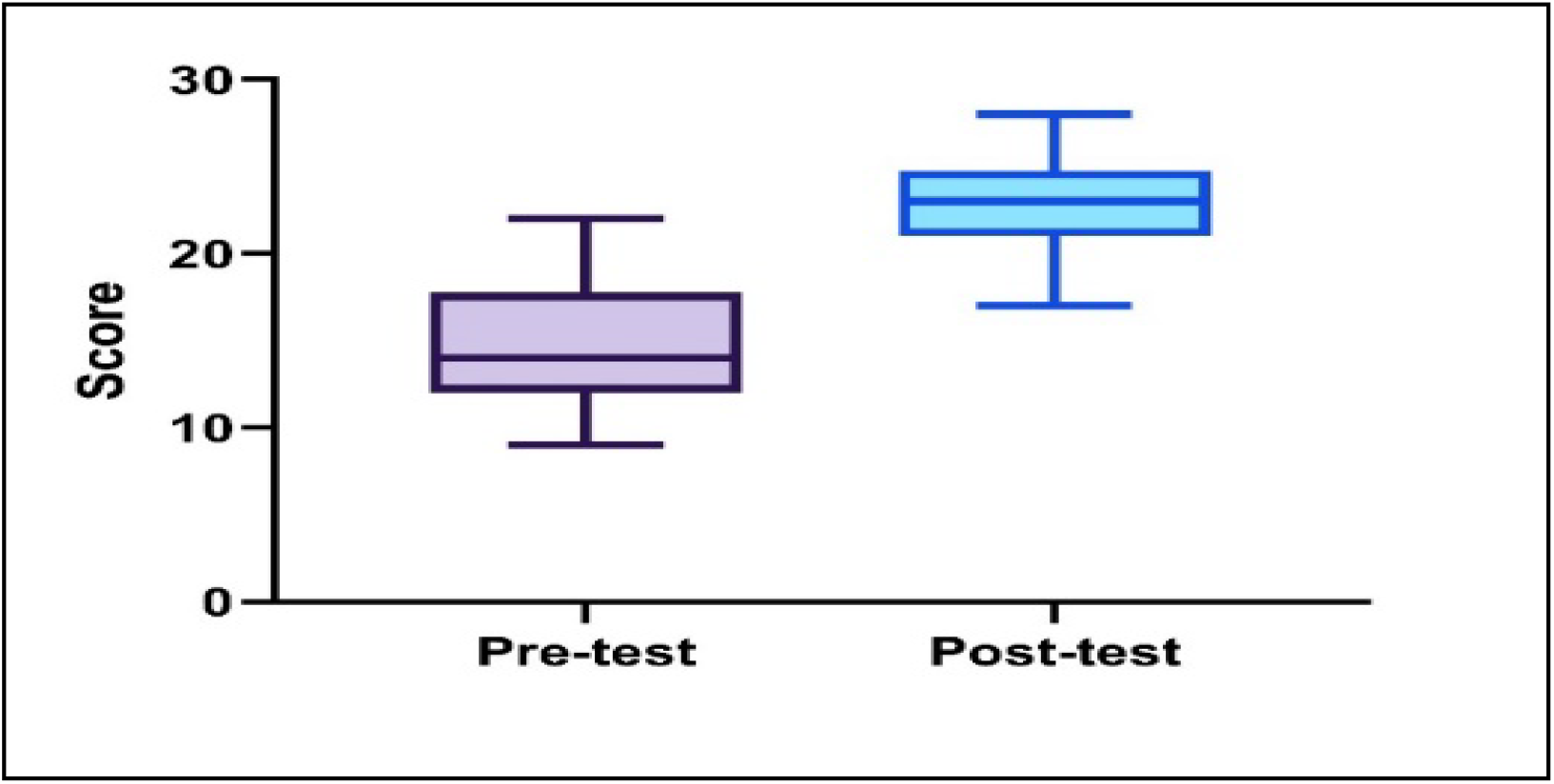
Pre-test and post-test score.

Analysis of the 11 questions used to assess caregivers’ knowledge and awareness about disease management displayed significant improvement (p < 0.05) in post-test scores after the training sessions regarding types of insulin, commonly used insulin regimens for T1DM management, potential complications, rapid-acting glucose foods, emergency kit contents, and situations requiring emergency referral, as detailed in Table 3.

**Table 2:** Themes and sub themes extracted from in-depth interview.

| Themes | Sub themes |
| --- | --- |
| Misunderstanding of treatment plan | a. Unable to titrate the correct dosage of insulin<br>b. Confusion about what type of food should and should not be given<br>c. In doubt when to seek medical emergency help. |
| Distress of giving insulin to the children too often | a. Child is often at unease taking insulin shots multiple times/day<br>b. Fear of pricking the child at multiple places in the body<br>c. Sometimes the insulin dosage is missing, and child has to suffer the consequence<br>d. Avoidance for social gatherings as the child must take insulin Infront of other people |
| Fear of getting pricked for glucose monitoring | a. Child often cries when sees a lancet for pricking, its painful site for a caregiver<br>b. As a caregiver, it's concerning to see child's finger tips turn black and painful |
| Financial burden for the care givers | a. Caregivers are unable to cater all the special needs of the child<br>b. Monthly treatment cost for the child is unbearable<br>c. Unable to pay the bills post hospitalization of child which happens often<br>d. Some special discounts or free insulin should be provided by the government for the benefit of the children who require lifelong insulin for survival |
| Psychological stress for care givers | a. Child is unable to play like other kids, gets exhausted often on physical activity.<br>b. We don't socialize much as special care for the child is required<br>c. We are scared sometimes what if something bad happens to a child at school<br>d. We didn't plan for another child thinking the same would happen to the second child. |

**Table 3:** The detailed Pre-test / Post-test analysis.

| Questions | Total marks | Criteria for scoring | Pre-test | Post-test | P Value |
| --- | --- | --- | --- | --- | --- |
| 1. Diagnostic criteria for T1DM | 1 | Mention the range of glucose levels to label it as T1DM. | 0.51±0.32 | 0.93±0.05 | < 0.001 |
| 2. Signs and symptoms of Hyperglycemia | 3 | Any 1 correct sign or symptom-1mark<br>Any 2 correct sign or symptom-2 marks<br>Any 3 correct sign and symptom-3 marks | 2.38±.067 | 2.51±0.56 | 0.083 |
| 3. Signs and symptoms of Hyperglycemia | 3 | Any 1 correct sign or symptom-1mark<br>Any 2 correct sign or symptom-2 marks<br>Any 3 correct sign and symptom-3 marks | 2.32±0.63 | 2.46±0.56 | 0.095 |
| 4.How do you monitor blood glucose levels at home | 1 | Name the device used to monitor the blood glucose level. | 0.97±0.17 | 1.00±0.00 | 0.159 |
| 5. Types of insulin | 3 | One correct type of insulin-1 mark<br>Two correct types of insulin-2 marks<br>Three correct types of insulin-3 marks | 1.65±0.62 | 2.47±0.59 | <0.001 |
| 6. Different sites for administration of insulin | 2 | One correct site- 1mark<br>Two correct sites-2 marks | 1.35±0.48 | 1.46±0.50 | 0.051 |
| 7. Name commonly used Insulin regimes for management of T1DM | 2 | Any one correct regime-1mark<br>Any two correct regime-2 marks | 0.44±0.50 | 1.72±0.45 | <0.001 |
| 8. What are the possible complications of T1DM? | 2 | Any one correct complication -1 mark<br>Any 2 correct complications -2 marks | 0.96±0.21 | 1.26±0.44 | <0.001 |
| 9. What do you do when you find out that your child has an episode of Hypoglycemia? | 2 | Any one correct step-1mark<br>Any two correct steps -2 marks | 0.97±0.17 | 1.07±0.26 | 0.018 |
| 10. Name some rapid acting glucose foods | 3 | Any one correct name-1mark<br>Any two correct names-2 marks<br>Any three correct names-3 marks | 0.99±0.21 | 1.79±0.78 | <0.001 |
| 11. Contents of Emergency kit. | 4 | Any one correct content - 1 mark<br>Any 2 correct contents-2 marks<br>Any 3 correct contents-3 marks<br>Any 4 correct contents-4 marks | 0.76±0.58 | 3.22±0.69 | <0.001 |
| 12. In which situations you refer your child to emergency? | 4 | Any one correct condition- 1 mark<br>Any 2 correct conditions - 2 marks<br>Any 3 correct conditions-3 marks<br>Any 4 correct conditons-4 marks | 0.91±0.81 | 3.26±0.64 | <0.001 |
| **Total score** | **30** |  | **14.26±3.34** | **23.17±2.28** | **<0.001** |

## DISCUSSION

Our study highlights those caregivers of children with T1DM experience misunderstandings about the treatment plan, distress from frequent insulin administration, and fear associated with glucose monitoring. They also face financial burdens and significant psychological stress. Importantly, the structured educational program significantly improved post-test questionnaire scores among caregivers, indicating enhanced understanding and management of the disease compared to pre-test scores.

Involving families in diabetes education programs is crucial for achieving good diabetes control in children.^7^ Studies show that caregivers who adopt coping strategies, such as acceptance and reliance on social support, experience less depression and foster better collaboration and adherence to diabetes management among their children.^8,9^ Pilot studies suggest that motivational interviewing (MI) and solution-focused brief therapy (SF) can effectively reduce HbA1c levels, alleviate fear of hypoglycemia, and improve quality of life (QoL).^10,11^

Managing T1DM requires a comprehensive and ongoing approach to care. A lack of consistent communication between healthcare providers and parents can diminish motivation to manage the child’s condition, further impacting QoL.^12^ This finding aligns with research by Dudley et al. (2014), which emphasizes that effective teamwork within the healthcare team significantly influences child health outcomes.^13^

Wigert et al. further stresses the importance of multidisciplinary teamwork in achieving better health outcomes in adolescents with T1DM. Clearly defined roles and responsibilities among team members are essential for improving patient outcomes.^14^ Healthcare providers should regularly engage with children and their families on a variety of topics, including school, diabetes camps, psychological issues, fears related to hypoglycemia, and future aspirations.^15,16^

By fostering awareness regarding disease management, including glucose monitoring, insulin titration, maintaining a healthy diet, and encouraging exercise, life-threatening complications from T1DM can be reduced significantly among these children.

One of the limitations of this research is the findings may not be generalizable, as it is a single center study. Self-reported data would introduce inaccuracies, and the short follow-up period also limits the assessment of long-term impacts. Variations in session delivery might affect consistency, and the focus, which was primarily on knowledge and practices, did not consider other outcomes, such as caregiver stress. Additionally, expert validation could be biased due to differing familiarity with caregivers’ needs.

## CONCLUSION

This study underscores the critical role of structured educational interventions in enhancing the knowledge and management practices of caregivers for children with type 1 diabetes mellitus (T1DM). It highlighted the diverse challenges faced by caregivers, including misunderstandings about treatment, psychological distress, and financial burdens. By addressing these issues, the educational program significantly improved caregivers’ knowledge and management skills, demonstrating its effectiveness.

The involvement of an interprofessional healthcare team was essential in delivering tailored support and fostering a collaborative approach to diabetes management. This study emphasizes the importance of ongoing education and support for caregivers, which is vital not only for improving disease outcomes but also for enhancing the overall well-being of both caregivers and children.

Ultimately, educational programs developed by interprofessional experts can effectively empower caregivers to better care for their children with T1DM. Future research should focus on multi-center studies to assess the long-term impact of such interventions and explore additional outcomes, such as caregiver stress and quality of life. By prioritizing caregiver education and support, we can better equip families to navigate the complexities of T1DM management, leading to improved health outcomes for children.

